# Behavioral and neuroimaging correlates of attentional biases to angry faces in individuals in remission from depression: a population-derived study

**DOI:** 10.1101/2023.03.13.532400

**Authors:** Jakub Nagrodzki, Luca Passamonti, Suzanne Schweizer, Jason Stretton, Ethan Knights, Richard Henson, Cam-CAN, Noham Wolpe

**Author notes:** Corresponding author Dr Jakub Nagrodzki, Department of Psychiatry, Herchel Smith Building for Brain & Mind Sciences, Forvie Site, Robinson Way, Cambridge CB2 0SZ, UK. https://www.cam-can.org/index.php?content=corpauth#14 (Cam-CAN).

## Abstract

**Background:** Depressed individuals show attentional biases in the processing of emotional stimuli, such as negative face expressions. Some of these biases persist in previously depressed individuals, but their mechanisms remain largely unknown.

**Methods:** A population-derived cohort (*n* = 134, 68 females; 21 - 92 years) was recruited by Cam-CAN. Functional MRI was acquired during a gender discrimination task, which used angry and neutral faces. Drift diffusion modelling (DDM) was used to investigate the latent components of the decision process, focusing on the effect of emotional valence. DDM parameters were correlated with activity in brain regions.

**Results:** 14% of participants reported a history of depression in remission. The best fitting DDM specified a different drift rate for angry and neutral faces. A slower drift rate for angry faces predicted depression in remission (OR 0.092, *p* = 0.048). This effect persisted after accounting for current depression symptoms and drift rate for neutral faces. Participants with a slower drift rate for angry faces demonstrated increased activations in the bilateral insula, bilateral inferior frontal gyrus and bilateral parietal cortex when viewing angry relative to neutral faces.

**Conclusions:** Our results suggest a persistent attentional bias in the processing of angry faces in individuals with depression in remission, over and above their current depressive symptoms. The imaging findings suggest that the slowing is associated with changes in areas involved in emotional regulation and evidence accumulation. Attentional biases in the processing of emotional information may reflect a trait, rather than state, in individuals with depression.

## INTRODUCTION

Depressed individuals process emotional stimuli differently from people without depression. For example, mounting evidence suggests that when looking at emotional face expressions, depressed individuals have an attentional bias towards negative faces (1–3). This attentional bias has also been reported in people who were previously depressed (4–6) and has been suggested to be causally involved in depression and risk of depression relapse (6,7).

One type of such negative stimuli are angry faces. Depressed individuals have been shown to have increased attentional engagement for angry, but not neutral, faces compared to non-depressed participants (2,8). Gilboa-Schechtman and colleagues (2004) offer a thorough account of why angry faces are particularly poignant stimuli in depression. Angry faces are perceived as threatening stimuli, which are ‘mood-eliciting’, rather than just mood-congruent. Rather than just signaling the emotional state of another person (like sad or fearful faces), these stimuli cause individuals to reappraise their own actions. It is therefore possible that previously depressed individuals will be highly sensitive to angry faces as threatening signals.

Experimentally, attentional biases can be demonstrated in a variety of ways – using reaction time (2,9), eye tracking (10), or task performance, such as task accuracy (11). The increased attention to the emotional component of the stimulus manifests as a performance enhancement when the emotional content is task-relevant, or a performance impairment when it is task-irrelevant (12). This may reflect the allocation of additional cognitive resources to process the emotional content (13). However, it remains unclear which latent cognitive process is influenced by attentional biases towards negative emotion (negativity attentional bias).

Sequential sampling models, such as the drift-diffusion model (DDM), can be used to investigate the underlying latent cognitive processes that explain changes in both reaction time and accuracy (14,15). The DDM assumes a stochastic accumulation of evidence until a threshold is crossed, at which point an individual commits to a decision (16). On this account, the attentional negativity bias in depression could be explained by a slower accumulation of evidence for negative emotional stimuli; or a more cautious behavior requiring more evidence to be accumulated to make decisions about negative stimuli.

Here, we sought to examine which latent cognitive variable can capture the attentional negativity bias in individuals with depression in remission. We further asked whether this latent variable would be able to distinguish individuals with or without a history of depression. Finally, we investigated what the neural correlates of this latent variable are. To this end, we combined an implicit emotion processing task, DDM and functional MRI (fMRI) to examine the behavioral and neural correlates of attentional bias for angry faces in individuals with depression in remission. We used a population-derived cohort of currently healthy participants across the adult lifespan. Participants performed an emotional face task, requiring them to discriminate the gender of angry and neutral faces (17).

We hypothesized that attentional negativity bias would be associated with a slower accumulation of evidence for negative emotional stimuli, in line with previous studies investigating how stimuli interfere with the decision process (18). We further hypothesized that this would be mediated by differences in brain activation in the limbic system, including the amygdala (12,19), and the attentional network, including the inferior frontal gyrus (20).

## METHODS AND MATERIALS

### Participants

A population-derived cohort of healthy adults (*n* = 136) was recruited as part of the third stage (‘CC280’) of the Cambridge Centre for Ageing and Neuroscience (Cam-CAN) (21,22). Exclusion criteria are described in Shafto et al. (2014) including: significant cognitive impairment (mini-mental state examination score of 24 or less), communication difficulties, significant medical problems (full list in Table 1 in Shafto et al., 2014), mobility problems, substance abuse, and MRI/MEG safety and comfort issues. Ethical approval was granted by the local ethics committee, Cambridgeshire 2 (now East of England – Cambridge Central) Research Ethics Committee (reference: 10/H0308/50). Written informed consent was obtained from all participants before commencing the study.

**Table 1.** Summary of the demographic details of participants included in the study. Number of participants (percentage, %) or median (interquartile range) is presented. N = number, GCSE = General Certificate of Secondary Education, HADS = Hospital Anxiety and Depression Scale, BFR = Benton Face Recognition score; Sig. = significance; ns = no statistically significant difference between the two groups at a level of p < 0.05; ** p < 0.01.

|  |  | Depression in remission | Never depressed | Total | Sig. |
| --- | --- | --- | --- | --- | --- |
| **N** |  | 19 | 115 | 134 |  |
| **Age** |  | 47.037<br>[38.109, 59.764] | 58.159<br>[40.635, 73.353] | 54.749<br>[39.184, 71.926] | ns |
| **Female** |  | 11 (58%) | 57 (50%) | 68 (51%) | ns |
| **Highest level of education** | *Before GCSE* | 2 (10%) | 4 (3%) | 6 (4%) | ns |
|  | *GCSE* | 1 (5%) | 7 (6%) | 8 (6%) |  |
|  | *A-Level* | 3 (15%) | 10 (9%) | 13 (10%) |  |
|  | *University* | 13 (68%) | 94 (82%) | 107 (80%) |  |
| **HADS depression** |  | 4 [3, 6.5] | 2 [1, 4] | 2 [1, 4] | ** |
| **BFR** |  | 23 [22, 25] | 23 [21, 25] | 23 [21, 25] | ns |

**Table 2.** (overleaf). Summary of fMRI results.

|  | MNI coordinates<br>of peak (mm) | AAL3 label | Cluster<br>size |
| --- | --- | --- | --- |
| **Contrast 1: Main effect of stimulus emotion (Angry – Neutral)** |  |  |  |
| Cluster 1 | 30 -90 3 | Right middle temporal gyrus<br>Right middle occipital gyrus<br>Right superior temporal gyrus<br>Right inferior occipital gyrus<br>Right superior occipital gyrus<br>Right cuneus | 1578 |
| Cluster 2 | -30 -93 6 | Left middle occipital gyrus<br>Left middle temporal gyrus<br>Left inferior occipital gyrus<br>Left superior temporal gyrus<br>Left superior occipital gyrus | 1252 |
| Cluster 3 | -42 -42 -18 | Left inferior temporal gyrus | 92 |
| Cluster 4 | 42 -39 -15 | Right fusiform gyrus | 100 |
| Cluster 5 | 54 33 6 | Right inferior frontal gyrus, triangular part<br>Right inferior frontal gyrus, orbital part | 184 |
| Cluster 6 | -48 30 0 | Left inferior frontal gyrus, triangular part | 102 |
| **Contrast 2: Main effect of stimulus emotion (Neutral – Angry)** |  |  |  |
| Cluster 1 | 21 30 -12 | Right superior frontal gyrus: medial orbital<br>Left superior frontal gyrus: medial orbital<br>Left rectus<br>Right rectus<br>Left inferior frontal gyrus, orbital part<br>Right ventral striatum<br>Left olfactory cortex<br>Left ventral striatum<br>Left posterior orbital gyrus<br>Left anterior cingulate cortex, pregenual<br>Right anterior cingulate cortex, subgenual<br>Right anterior orbital gyrus<br>Left anterior cingulate cortex, subgenual<br>Right medial orbital gyrus<br>Right posterior orbital gyrus<br>Left medial orbital gyrus | 787 |
| Cluster 2 | -6 -3 54 | Left supplementary motor area | 81 |
| Cluster 3 | 60 -39 -18 | Right inferior temporal gyrus | 60 |
| **Contrast 3: Main effect of drift rate for angry faces (Angry – Neutral)** |  |  |  |
| **No significant clusters** |  |  |  |
| **Contrast 4: Main effect of drift rate for angry faces (Neutral – Angry)** |  |  |  |
| Cluster 1 | -42 12 6 | Left insula<br>Left inferior frontal gyrus, opercular part | 96 |
| Cluster 2 | 36 21 9 | Right insula<br>Right inferior frontal gyrus, opercular part | 146 |
| Cluster 3 | -39 -48 42 | Left inferior parietal gyrus | 93 |
| Cluster 4 | 48 -33 39 | Right supramarginal gyrus | 301 |

In the initial interview process, participants were asked whether they had ever been diagnosed and treated for depression, and if so when. Demographic information, such as level of education, was also obtained. One participant who declared they had depression diagnosed in the same year was excluded from further analysis. Participants were also administered the Benton Test of Facial Recognition (Levin et al., 1975).

### Behavioral task

Participants undertook a gender discrimination task of emotional faces, requiring them to identify the gender of a face showing an angry or neutral face expression (Fig. 1) (Passamonti et al., 2008). Participants were instructed to respond as quickly and as accurately as possible. Instructions and visual stimuli were back-projected onto a screen viewed through a mirror mounted on the MRI head coil. The stimuli consisted of 60 faces (30 identities; equal numbers of male and female identities), which displayed either an angry or a neutral face expression.

**Figure 1.**
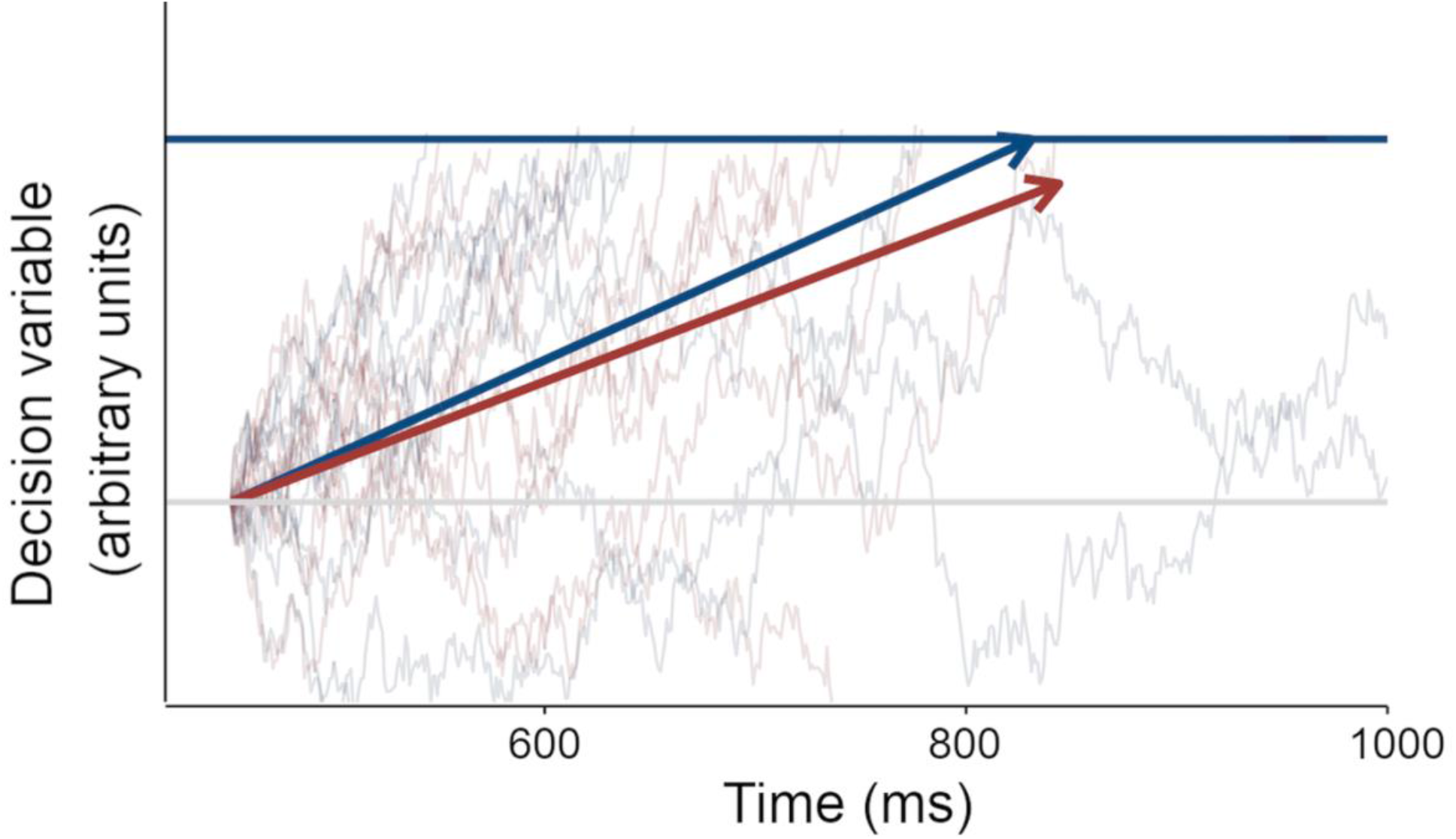
Simulation of the drift diffusion process, based on the drift rate for neutral (blue) and angry (red) faces. The grey horizontal line represents the starting point of the drift process, the blue horizontal line represents the decision threshold. Twenty trials were simulated for illustration. The arrows represent the mean of all traces.

The experiment consisted of 24 blocks (12 angry and 12 neutral), each lasting 21 s. Each block consisted of six face trials, which were pseudo-randomly interleaved with six null events (central cross), such that there were no more than three consecutive trials of the same type (face or null). During each face trial, a face was presented for 1000 ms, followed by a fixation point (central cross) for 750 ms. During each null event, the central cross was displayed for 1750 ms. Participants responded with a button press to each face to indicate whether it was male or female.

The principal measures for each trial were reaction time (RT) and accuracy, i.e., whether the participant correctly identified the gender of the face stimulus. One participant responded incorrectly to >40% of the trials and was therefore excluded from further analysis. The data from the remaining participants (*n* = 134) were used for the behavioral and imaging analyses.

### Drift-diffusion modelling

To investigate the latent cognitive components of the decision-making process contributing to the variability in task performance and reaction time, DDMs were fit using the HDDM toolbox for Python, v0.9.2 (23). Bayesian hierarchical model fitting was used to estimate each participant’s model parameter, as drawn from a group distribution. In line with previous studies in the field, ultra-fast RTs shorter than 250 ms were removed (Wiecki et al., 2013). Overall, 3.3% of the total trials were excluded in this way. Other default parameters were used as suggested (Wiecki et al., 2013).

For our main analyses, we considered accuracy coding models, as we sought to explain both accuracy and RT in the task, both of which have been shown to be affected by emotion (2,9,11). For completeness, we also report the parameters of stimulus coding models (Supplementary Material). The model parameters included the drift rate (v), boundary separation (a), and non-decision time (t). We employed a pragmatic restriction on model complexity for interpretability, with just one model parameter dependent on stimulus emotion at a time. We fit four different models initially, with one model parameter dependent on stimulus emotion (angry vs. neutral) per model:

- model 1 (the null model) with all parameters independent of stimulus emotion;
- model 2 with drift rate (v) dependent on stimulus emotion;
- model 3 with boundary separation (a) dependent on stimulus emotion;
- and model 4 with non-decision time (t) dependent on stimulus emotion.

Each model was estimated 5000 times, discarding the first 1000 samples to minimize the effect of initial values on posterior inference, and with a thinning factor = 2 to reduce autocorrelations. Model convergence was assessed with visual inspection of the posterior plots of the samples. The deviance information criterion (DIC) was calculated for each model and the model with the lowest DIC was selected for further analysis.

The best fitting model was run with 5 chains, 10,000 samples each. The first 2000 samples were discarded as burn-in and a thinning factor of 5 was used. Posterior convergence was assessed with the potential scale reduction statistic 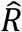 (< 1.1 for all parameters) and with inspection of posterior plots. One value of each model parameter was calculated per participant as the mean from each of the 8000 resulting model parameter estimates, for group comparison (see below).

Group comparisons for demographics and behavioral data (depressed in remission vs. never depressed) were performed with SciPy (24), using: the t-test, the Mann-Whitney U test for comparing ranks where normality assumption was violated, or the Chi-squared test for proportions, while the stats package was used for logistic regression in R (25).

### Functional brain imaging acquisition and analyses

While participants performed the behavioral task, fMRI data were acquired on a 3T Siemens TIM Trio System, employing a 32-channel head coil, using T2*- weighted contrast from a Gradient-Echo Echo-Planar Imaging (EPI) sequence. A total of 381 volumes were obtained per participant, each containing 32 axial slices (in descending order). Slice thickness was 3.7 mm with an interslice gap of 20%; TR = 2 s; TE = 30 ms; flip angle = 78 degrees; FOV = 192 × 192 mm; voxel-size = 3 × 3 × 4.44 mm). Total acquisition time was 12 mins and 27 s.

The fMRI data were preprocessed and analyzed using SPM12 (https://www.fil.ion.ucl.ac.uk/spm/software/spm12/) in MATLAB (Mathworks, MA). Details of the Cam-CAN preprocessing pipelines have been described at length previously (22). In short, data were unwarped using field-map images, realigned to correct for motion, slice-time corrected, and co-registered to each participant’s T1-weighted image. The normalization parameters from applying DARTEL to the structural image (26) were then applied to warp functional images into MNI space. The scans were smoothed with an 8 mm isotropic Gaussian kernel.

For each participant, a general linear model (GLM) was fit to the fMRI timeseries in each voxel, which included regressors formed by convolving the estimated neural activity for each condition with a canonical hemodynamic response function. Neural activity for each trial was modelled as a boxcar with duration equal to the reaction time for that trial. Three conditions were defined: the two experimental conditions (angry, neutral), as well as an ‘invalid’ condition for trials with no button press, or ultrafast trials with RT < 250 ms. Six motion parameters were added to account for motion-related variability, resulting in nine regressors in total. A high-pass filter with a cut-off of 128 s was applied, and an autoregressive model was used to estimate autocorrelation in the data, and inverted to pre-whiten the data and model.

The difference in the estimated parameters for angry versus neutral faces was calculated for every voxel to produce an image of this contrast. These images were entered into two group-level GLMs. This first GLM included participant group (depressed in remission or never depressed). This enabled us to compare whether the difference in activation between neutral and angry faces was further different for each group. The second group-level GLM included the drift rate for neutral faces and the drift rate for angry faces as regressors, enabling us to identify how the activity in the angry vs. neutral condition correlated with the drift rate for angry faces, above and beyond a correlation with drift rate for neutral faces. In a final control analysis, we included several possible confound variables, in addition to drift rates, including age, sex, education, HADS depression and Benton face recognition score (see Supplementary Material).

To control for family-wise error (FWE) at p < 0.05, random field theory was used for cluster-level inference, given an initial cluster-forming threshold of *p* < 0.001, uncorrected. An explicit grey matter mask was obtained from the DARTEL template and applied to the results. Mapping of clusters to anatomical areas was performed using the Automated Anatomical Labelling atlas 3 (AAL3) extension for SPM12 (27). Labels that overlap with at least 5% of voxels in a cluster were reported.

## RESULTS

### Participants

A summary of the demographic details of participants included in the study is reported in Figure 2 for each group (DIR vs. never depressed).

**Figure 2.**
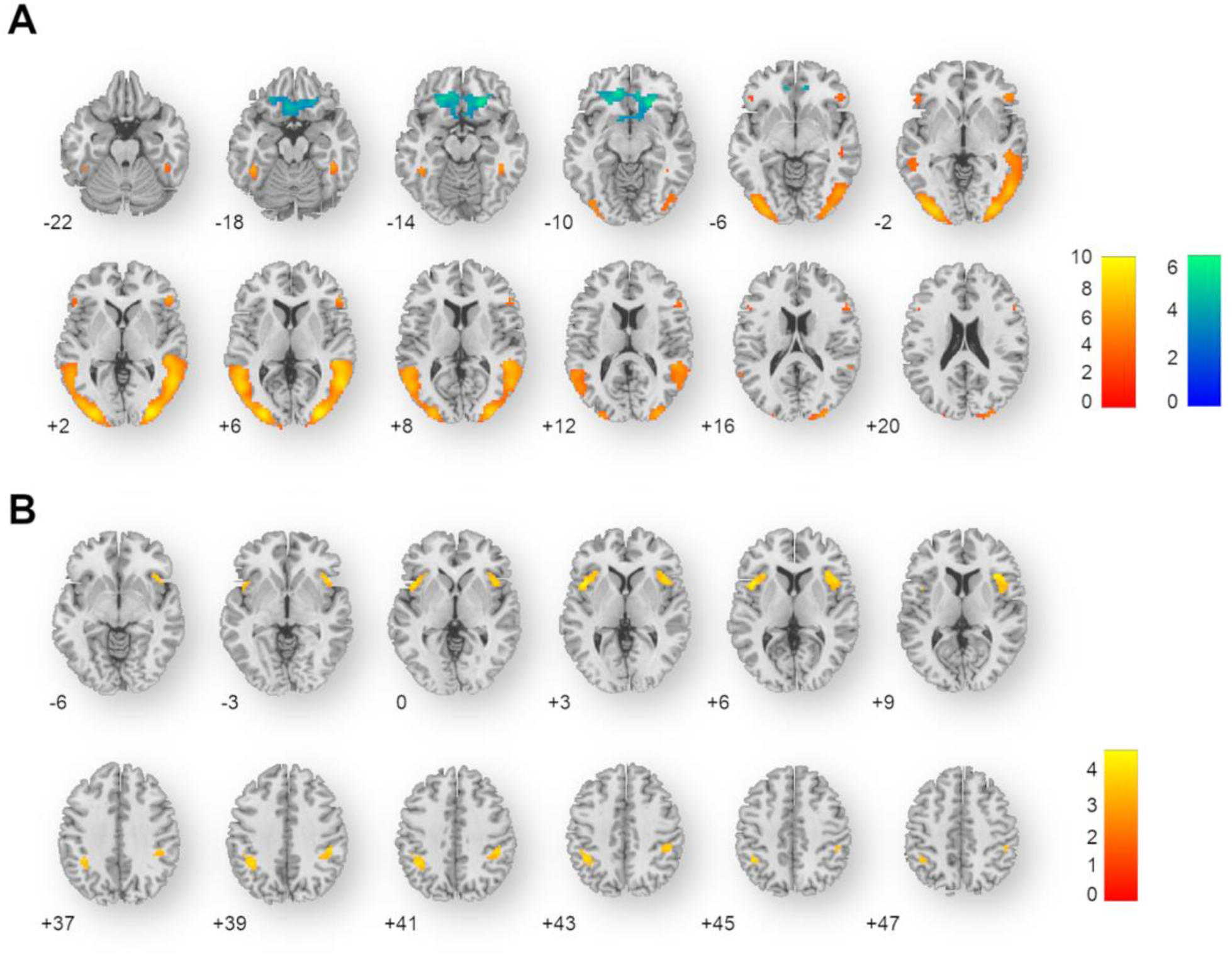
Summary of fMRI results. Axial slices are shown, numbers indicate z coordinates of the slice. A) Axial slices showing areas of increased (warm color scale) and decreased activation in the [angry – neutral] contrast in all participants. B) Axial slices showing areas of increased activation with decreasing drift rate for angry faces in the [angry – neutral] contrast in all participants.

The two groups differed only on HADS depression scores, with the DIR group having higher HADS depression scores (median = 4, interquartile range (IQR) = [3, 6.5]) than the never depressed group (median = 2, IQR = [1, 4]; *U =* 663.5, *p* = 0.003). There was no statistically significant difference between the two groups in age (DIR: median = 47.037, IQR = [38.109, 59.764], never depressed: median = 58.159, IQR = [40.635, 73.353]; *U =* 845), proportion of females (*X*^*2*^(1, *N* = 134) = 0.181), level of education (*X*^*2*^(3, *N* = 134) = 3.036) or Benton Face Recognition score (median = 23, IQR = [22, 25]; median = 23, IQR = [21, 25]; *U = 1039*), all *p*s > 0.06.

### Emotional processing and drift-diffusion modelling

An analysis of reaction time revealed no difference between angry and neutral stimuli, but did show a significant difference in accuracy, with increased accuracy in the neutral condition (Supplementary Material).

We fit a set of DDMs to the behavioral data, varying the fixed and free parameters (see Methods). Model DICs (goodness of fit) were as follows:

- model 1 (the null model) DIC = 7146;
- model 2 with drift rate (v) dependent on stimulus emotion DIC = 6970;
- model 3 with boundary separation (a) dependent on stimulus emotion DIC = 7104;
- model 4 with non-decision time (t) dependent on stimulus emotion DIC = 7146.

The model with the lowest DIC was selected for further analysis. This model included both the boundary separation ‘a’ and non-decision time ‘t’ fixed across both conditions and, but importantly, a different drift rate for each of the two experimental conditions, namely angry and neutral. Across all participants, the drift rate was significantly slower for angry faces (mean = 1.899; standard deviation = 0.557) than for neutral faces (2.215; 0.528), *t*(265) *=* -4.748, *p* = 3e-06. Together, the results indicate that participants accumulated evidence about the stimulus gender more slowly in the angry condition compared to the neutral condition, suggesting an attentional negativity bias.

We tested whether individual differences in attentional negativity bias as quantified by the drift rates were associated with a history of depression. A logistic regression was performed, with drift rates for angry and neutral faces predicting patient depression history. We further included HADS depression as a covariate to rule out current depressive symptoms and age, given the large age range in the study, as well as the higher cumulative likelihood of having been diagnosed with depression with increasing age.

The logistic regression model was statistically significant, *X*^*2*^(5, *N* = 134) = 17.334, *p* = 0.002. The model explained 21.7% (Nagelkerke *R*^2^) of the variance in incidence of depression in remission. Slower drift rate for angry faces significantly increased the likelihood of belonging to the DIR group (OR 0.091, 95% CI [0.007; 0.866], *p* = 0.047). The drift rate for neutral faces was not significantly associated with depression history (OR 5.002, 95% CI [0.583; 50.754], *p* = 0.152). In addition, an increase in age by 1 year decreased the likelihood of belonging to the DIR group by 4.9% (95% CI [1.2%; 8.4%], *p* = 0.011), whereas an increase in the HADS depression score by 1 point increased the likelihood of belonging to the DIR group by 28.6% (95% CI [8.5%; 54%], *p* = 0.004). These findings suggest that a slower drift rate for angry faces is a persistent finding in individuals who are in remission from depression, even after accounting for their age and any residual depressive symptoms. This effect persisted after additionally accounting for all other demographic factors (sex, education, Benton face recognition score; see Supplementary Material). In a further exploratory control analysis, medication status had no effect on the drift rate for angry faces (Supplementary Material).

### Functional brain imaging results

The results of the fMRI analyses are summarized in Figure 4. Viewing angry compared to neutral faces showed widespread increased activity in the bilateral occipital and temporal areas, including the right fusiform gyrus, as well as in the inferior frontal gyri. Decreased activity was observed in a large cluster encompassing bilateral orbitofrontal cortex and anterior cingulate cortex, and two more localized clusters encompassing the left supplementary motor area and the right inferior temporal gyrus. There were no significant differences in this activity patterns between DIR and never depressed groups.

We then examined the influence of drift rate for angry faces on activations related to viewing angry vs. neutral face expressions across all participants. Participants with a slower drift rate for angry faces demonstrated increased activations in the bilateral insulae, bilateral inferior frontal gyri, and bilateral parietal cortex when viewing angry compared to neutral faces. Including the remaining demographic factors (age, sex, education, Benton face recognition score, HADS depression) did not change the results (see Supplementary Material).

## DISCUSSION

Our study combines computational modelling of behavior with neuroimaging in a population-derived cohort to examine differences in negativity attentional bias. Our results show that individuals with depression in remission show slower rate of evidence accumulation for negative (angry) emotional stimuli, above and beyond their current depressive symptoms. In addition, across all participants a slower rate of evidence accumulation for angry faces is associated with greater activity in bilateral insula, bilateral inferior frontal gyrus, and bilateral parietal cortex.

We found that the drift rate (speed of evidence accumulation) was sensitive to the emotional expression of the faces, which in our task was irrelevant to the task itself (the face gender). Across participants, evidence accumulation for the gender decision was slower for angry compared to neutral faces. This finding corroborates previous studies, which consistently show that participants spend longer looking at threatening face expressions (e.g., Belopolsky et al., 2011) and that the presentation of emotional stimuli leads to slower and less accurate responses when the emotional content acts as a distractor (3). This may reflect the need to allocate additional cognitive resources to the processing of emotional content, over and above those required to complete the task at-hand (13). We identified a cognitive process (slower evidence accumulation rate) underlying this observational evidence (increased attention to emotional stimuli).

Using DDM to estimate this underlying cognitive process, the principal result of our study is that a slower evidence accumulation rate for angry stimuli is associated with depression in remission. This is consistent with previous research demonstrating that biased processing of emotional information persists in previously depressed (currently in remission) individuals (1,2). Importantly, we found that this effect is above and beyond their current depressive symptoms and age, despite both higher residual depressive symptoms and increasing age being associated with a greater cumulative risk of having suffered a depressive episode in the past. One possibility is that the slower evidence accumulation rate for angry stimuli reflects an individual trait, rather than a state that is only present during a depressive episode (6).

This result raises an interesting possible clinical application. Attentional biases in stimulus processing have been suggested to be causally involved in depression and risk of depression relapse (6,7). Individual differences in measures of negativity attentional bias (such as drift rate) may therefore identify individuals who are at risk of depression relapse. In other words, previously depressed individuals showing persistently slower drift rate for negative (e.g., angry) emotional stimuli may, speculatively, be at higher risk of relapse and warrant preventative treatment (28). Further longitudinal research is required to test this clinically relevant hypothesis.

Our neuroimaging data show that, across all participants, a slower drift rate for angry faces is related to greater activity in the bilateral insulae, inferior frontal gyri and parietal cortex when viewing angry faces. The insula is part of the “rich club” of highly connected brain regions (Dai et al., 2015; Harriger et al., 2012) and is frequently reported in imaging studies in healthy individuals and in people with mental health disorders, including depression and other mood disorders (31–33). It also forms part of the salience network, where it is suggested to prioritize salient information for neural processing (34,35), and is therefore likely involved in attentional biases towards negative emotional stimuli.

The inferior frontal gyrus (IFG) has been typically implicated in behavioral inhibition (right IFG, Aron et al., 2003; Duann et al., 2009), and its activity in depression is thought to reflect response to treatment (Dai et al., 2020; Gorka et al., 2019; Marwood et al., 2018). Moreover, while some studies have also reported changes in the activity of the parietal cortex in major depressive disorder (41,42), it is more likely that our neuroimaging finding is related more to its role in evidence accumulation. Human (43–45) and non-human primate (46–49) research has shown that the parietal cortex is the site of evidence accumulation in decision-making tasks. Given that our principal result refers to a measure of speed of evidence accumulation, the association between this measure and parietal activity is not surprising. Together, the areas identified in our study could form a network, whereby the prefrontal cortex modulates parietal-mediated evidence accumulation (50,51), informed by emotional salience or valence signaled by the insula (35).

Our study has several limitations. First, we rely on self-report of depression history, rather than clinical records. The accuracy of self-reported depression history is clearly limited, mainly in underestimating depression prevalence (52). Interestingly, in our study increasing age predicted a lower risk of having depression in remission, suggesting either poor recall of historical diagnoses or a generational effect of lower rate of incidence, diagnosis or reporting in the older participants. However, the prevalence of prior depression diagnosis in our population-derived sample (14.2%) was similar to that in population-representative studies (53). Moreover, an inaccurate recall of depression history would lead to incorrect group classification in our study (for example, wrongly classifying a previously depressed individual to the never depressed group), thereby making it more difficult to detect a significant group effect. Second, our study is cross-sectional in nature and does not provide follow-up clinical information, for example recurrence of depression. Longitudinal studies are paramount for establishing the ability of DDM measures in a similar task to predict relapse risk. Third, as a population-derived study, only 19 individuals reported a previous diagnosis of depression, leading to a small group size. Finally, depression in remission as defined here yields a highly heterogenous group in terms of severity, clinical characteristics, treatment received and psychosocial factors. One could argue that these make any group differences even more compelling.

In summary, we find that a negativity attentional bias, namely slower drift rate for angry faces, predicts being previously depressed and currently in remission, and is related to differences in brain activity in regions related to emotion processing. These findings have implications to understanding cognitive biases in, and vulnerability to, depression.

## ACKNOWLEDGEMENTS

We are grateful to the Cam-CAN respondents and their primary care teams in Cambridge for their participation in this study. We also thank Dr Margaret Westwater for her advice on statistical analysis.

Cam-CAN research was supported by the Biotechnology and Biological Sciences Research Council (BB/H008217/1). JN was supported by an Academic Foundation Programme and BIRAX Ageing Travel Grant. NW was supported by an Israel Science Foundation Personal Research Grant 1603/22 and a National Institute of Health Research (NIHR) Academic Clinical Fellowship (ACF-2019-14-013). SS was supported by a Henry Wellcome Fellowship (209127/A/17/Z). RH was supported by MRC Programme Grant SUAG/086 G116768. For the purpose of open access, the author has applied a Creative Commons Attribution (CC BY) licence to any Author Accepted Manuscript version arising from this submission.

We thank the Cam-CAN project principal personnel: Lorraine K Tyler, Carol Brayne, Edward T Bullmore, Andrew C Calder, Rhodri Cusack, Tim Dalgleish, John Duncan, Richard N Henson, Fiona E Matthews, William D Marslen-Wilson, James B Rowe, Meredith A Shafto; Research Associates: Karen Campbell, Teresa Cheung, Simon Davis, Linda Geerligs, Rogier Kievit, Anna McCarrey, Abdur Mustafa, Darren Price, David Samu, Jason R Taylor, Matthias Treder, Kamen A Tsvetanov, Janna van Belle, Nitin Williams, Daniel Mitchell, Simon Fisher, Else Eising, Ethan Knights; Research Assistants: Lauren Bates, Tina Emery, Sharon Erzinçlioglu, Andrew Gadie, Sofia Gerbase, Stanimira Georgieva, Claire Hanley, Beth Parkin, David Troy; Affiliated Personnel: Tibor Auer, Marta Correia, Lu Gao, Emma Green, Rafael Henriques; Research Interviewers: Jodie Allen, Gillian Amery, Liana Amunts, Anne Barcroft, Amanda Castle, Cheryl Dias, Jonathan Dowrick, Melissa Fair, Hayley Fisher, Anna Goulding, Adarsh Grewal, Geoff Hale, Andrew Hilton, Frances Johnson, Patricia Johnston, Thea Kavanagh-Williamson, Magdalena Kwasniewska, Alison McMinn, Kim Norman, Jessica Penrose, Fiona Roby, Diane Rowland, John Sargeant, Maggie Squire, Beth Stevens, Aldabra Stoddart, Cheryl Stone, Tracy Thompson, Ozlem Yazlik; and administrative staff: Dan Barnes, Marie Dixon, Jaya Hillman, Joanne Mitchell, Laura Villis.

## Data availability and analysis code

All behavioral and imaging data are available through the Cam-CAN data repository (https://camcan-archive.mrc-cbu.cam.ac.uk/dataaccess). All analysis scripts are available via a GitHub repository under the MIT license agreement (https://github.com/jaknag/angry-faces-prev-depressed).

## DISCLOSURES

JN has nothing to disclose. LP has nothing to disclose. SS has nothing to disclose. JS has nothing to disclose. EK has nothing to disclose. RH has nothing to disclose. NW has nothing to disclose.

## Notes

https://camcan-archive.mrc-cbu.cam.ac.uk/dataaccess

### Competing Interest Statement

The authors have declared no competing interest.

