## Supplemental material for "Behavioral and neuroimaging correlates of attentional biases to angry faces in individuals in remission from depression: a population-derived study"

### SUPPLEMENTARY MATERIAL

#### Reaction time and accuracy analysis

For completeness, we report the results of the analysis of standard behavioural data (reaction time and accuracy).

Across all participants, the reaction time was not significantly different between angry faces (median = 0.741; IQR = [0.642; 0.891]) and neutral faces (0.744; [0.646; 0.887]), *U =* 9.96e07, *p* = 0.317. The accuracy of responses was significantly dependent on the condition (angry vs neutral) *X^2^*(1**,** *N* = 28274) = 308, *p* = 7.453e-69. The DDM with accuracy coding is sensitive to both changes in accuracy and reaction time and was employed in all further analyses.

#### DDM with stimulus coding

For completeness, we used DDM with stimulus coding to exclude the role of stimulus identity on our results. Using this method, response boundaries are defined not as ‘correct’ and ‘incorrect’ (accuracy coding), but rather as the two options presented to the participant, namely ‘male’ or ‘female’. A bias parameter (z) could then be added to investigate stimulus-related bias on the evidence accumulation process.

Drift-diffusion modelling (DDM) was performed using the same methodology as described in the ‘Drift-diffusion modelling’ section of the Methods in the main text, with the following modifications. The model parameters included the drift rate (v), boundary separation (a), and non-decision time (t), as well as the bias parameter (z). We fitted 29 different models, defined by a pragmatic restriction on model complexity, such that a maximum of one model parameter can depend on the stimulus gender, and a maximum of one model parameter can depend on the stimulus emotion in each model. This way, 29 models were specified (Fig. S1).

Goodness of fit was again estimated using the DIC. The best fitting model (Model 24) did not specify that the bias depended either on the stimulus gender or stimulus emotion. Similar to the accuracy coding model in the main text, drift rate (here towards ‘male or ‘female’ decision) depended on stimulus emotion, as well as on stimulus gender. Importantly, the winning model did not include a bias parameter that depended on the emotion or stimulus gender.

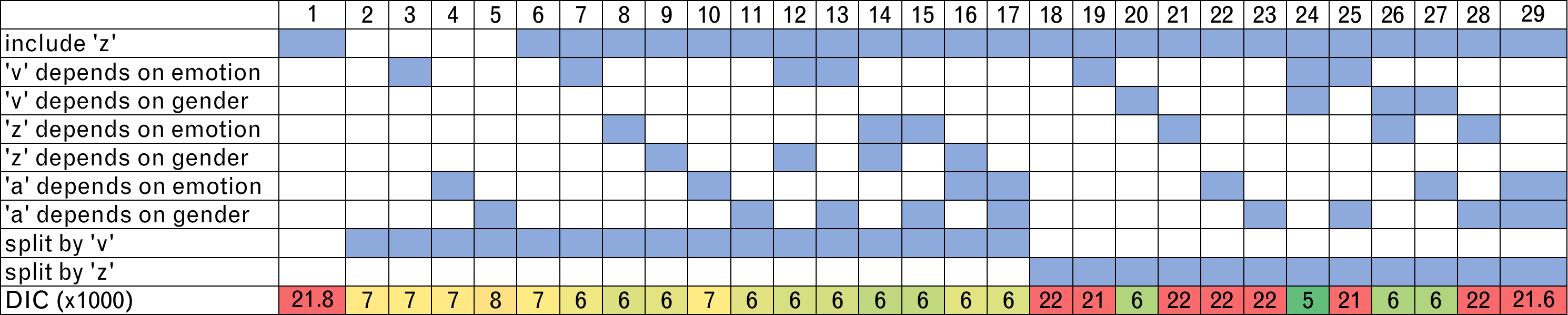

###### Figure S1. Model space and goodness of fit by DIC for drift-diffusion models with accuracy coding.

#### Control analysis of behavioural data

In order to exclude the contribution of current antidepressant therapy to the individual value of drift rate for angry faces in the depressed group, we conducted a multiple linear regression. The drift rate for angry faces was significantly associated with age (*β* = -0.019, *p* = 1.12e-14), but not with HADS depression (*β* = -0.011, *p* = 0.445) or the use of antidepressants (*β* = -0.043, *p* = 0.797).

We further explored the contribution of demographic cofounders to the logistic regression model by conducting a control analysis of the relationship between group (DIR vs never depressed) and the following covariates: the drift rate for angry stimuli, the drift rate for neutral stimuli, HADS depression, age, sex, highest educational qualification, Benton face recognition score.

The logistic regression model was statistically significant, *X^2^*(8, *N* = 134) = 23.922, *p* = 0.001. The model explained 29.3% (Nagelkerke *R*^2^) of the variance in incidence of depression in remission and correctly classified 88.1% of cases. Holding all other variables constant, it was found that a slower drift rate for angry faces significantly increases the likelihood of belonging to the DIR group (OR 0.058, 95% CI [3.547e-03; 0.639]). An increase in the HADS depression score by 1 point increased the likelihood of belonging to the DIR group by 34% (95% CI [12%; 64%]), whereas an increase in the BFR by 1 point increased the likelihood of belonging to the DIR group by 37% (95% CI [3.4%, 90%]). Age did not significantly contribute to the model (OR 0.961, 95% CI [9.202e-01; 1.001] and neither did the drift rate for neutral faces (9.790, [0.103; 117.302]), participant sex (0.759, [0.225; 2.460] or highest level of education (0.543 [0.260; 1.156].

#### Control fMRI analysis

Participant-level contrast images were constructed as described in Methods. These images were then entered in the control analysis, in which the group-level GLM included: drift rate for angry faces, drift rate for neutral faces, age, sex, education, HADS depression and BFR as regressors. The remainder of the analysis was conducted as described in Methods. The resultant clusters were identical to the ones obtained by way of the analysis reported in the main paper.
